# Ras/ERK-signalling promotes tRNA synthesis and growth via the RNA polymerase III repressor Maf1 in *Drosophila*

**DOI:** 10.1101/092114

**Authors:** Shrivani Sriskanthadevan-Pirahas, Rujuta Deshpande, Byoungchun Lee, Savraj S. Grewal

## Abstract

The small G-protein Ras is a conserved regulator of cell and tissue growth. These effects of Ras are mediated largely through activation of a canonical RAF-MEK-ERK kinase cascade. An important challenge is to identify how this Ras/ERK pathway alters cellular metabolism to drive growth. Here we report on stimulation of RNA polymerase III (Pol III)-mediated tRNA synthesis as a growth effector of Ras/ERK signalling in *Drosophila*. We find that activation of Ras/ERK signalling promotes tRNA synthesis both in vivo and in cultured *Drosophila* S2 cells. We also show that Pol III function is required for Ras/ERK signalling to drive proliferation in both epithelial and stem cells in *Drosophila* tissues. We find that the transcription factor Myc is required but not sufficient for Ras-mediated stimulation of tRNA synthesis. Instead we show that the main way that Ras promotes Pol III function and tRNA synthesis is by inhibiting the nuclear localization and function of the Pol III repressor Maf1. We propose that inhibition of Maf1 and stimulation of tRNA synthesis is one way by which Ras signalling enhances protein synthesis to promote cell and tissue growth.

## INTRODUCTION

The Ras small G-protein is one of the key conserved regulators of cell growth and proliferation. Over three decades of research have defined the textbook model of how Ras is activated by growth factors to stimulate a core RAF kinase, MEK (Mitogen-activated protein kinase kinase) and ERK (Extracellular signal-regulated kinase) signalling cascade. Work in model organisms such as *Drosophila, C elegans* and mouse has shown how this Ras/ERK pathway coordinates tissue growth and patterning to control organ size during development and homeostatic growth in adults.

Given its central role in development it is not surprising that defects in Ras signalling contribute to disease. Most notably, activating mutations in Ras and RAF occur in a large percentage of cancers, and lead to hyper-activation of ERK, which drives tumour formation in both epithelial and stem cells (Pylayeva-Gupta et al, 2011). Ras pathway mutations are also seen in several genetic developmental disorders - described collectively as RASopathies -often characterized by abnormal growth(Rauen, 2013). Understanding how Ras promotes cell proliferation and tissue growth is therefore an important concern in biology.

*Drosophila* has been a powerful model system to understand the biological roles of Ras signalling. In flies, Ras functions downstream of epidermal growth factor (EGF) and activation of its tyrosine kinase receptor (the EGFR). A series of genetic screens initiated in the late 80’s were pivotal in defining the canonical EGFR/Ras/ERK pathway (Karim et al, 1996; Rubin et al, 1997; Wassarman et al, 1995). Extensive studies since then have established when, where and how the pathway is activated during the fly life cycle to control development. This work has emphasized the importance of Ras signalling in the control of cell growth and proliferation (e.g. (Asha et al, 2003; Jiang & Edgar, 2009; Ninov et al, 2009; Parker, 2006; Read et al, 2009). Notably, during larval development Ras/ERK promotes EGFR-mediated cell proliferation and tissue growth in epithelial organs such as the imaginal discs, which eventually give rise to adult structures such as the legs, wings and eyes (Halfar et al, 2001; Karim & Rubin, 1998; Nagaraj et al, 1999; Prober & Edgar, 2000; Zecca & Struhl, 2002). In addition, in the adult the EGFR/Ras/ERK signalling controls proliferation of stem cell populations to maintain homeostasis and promote regenerative growth (Biteau & Jasper, 2011; Buchon et al, 2010; Castanieto et al, 2014; Jiang et al, 2011; Xu et al, 2011).

How does Ras mediate these effects on cell and tissue growth? Most work on this area has focused on transcriptional effects of Ras signalling. Work in *Drosophila* has identified several transcription factors that are targeted by ERK such as *fos, capicúa*, and *pointed*, and that regulate growth (Baonza et al, 2002; Biteau & Jasper, 2011; Jin et al, 2015; Tseng et al, 2007). Ras signalling has also been shown to crosstalk with other transcriptional regulators of growth such as the *hippo/yorkie* pathway and *dMyc* (Herranz et al, 2012a; Herranz et al, 2012b; Prober & Edgar, 2000; Prober & Edgar, 2002; Reddy & Irvine, 2013; Ren et al, 2013). These transcriptional effects control expression of metabolic and cell cycle genes important for growth(Baonza et al, 2002; Jin et al, 2015). Less is known, however, about how Ras/ERK may regulate mRNA translation to drive growth. The prevailing view, arising mostly from mammalian tissue culture experiments, is that ERK controls protein synthesis by stimulating the activity of translation initiation factors(Roux & Topisirovic, 2012). In particular, these effects are mediated via two ERK effector families - the MNK (MAP kinase-interacting serine/threonine-protein kinase) and RSK fribosomal s6 kinase) kinases (Roux et al, 2007; Roux & Topisirovic, 2012; Waskiewicz et al, 1999). These kinases are important for cellular transformation and tumour growth in mammalian cells (Romeo et al, 2013; Romeo & Roux, 2011; Ueda et al, 2010; Ueda et al, 2004). However, MNK and RSK mutants in mice and *Drosophila* have little growth or developmental phenotypes, and mouse MNK mutant cells show no alterations in protein synthesis (Arquier et al, 2005; Dumont et al, 2005; Reiling et al, 2005; Ueda et al, 2010; Ueda et al, 2004). These findings suggest Ras uses additional mechanisms to control translation and growth in vivo during animal development.

In this paper we report that the Ras/ERK pathway can stimulate RNA polymerase III-dependent tRNA synthesis. We find that these effects are required for Ras to drive proliferation in both epithelial and stem cells. Finally, we show that the main way that ERK promotes tRNA synthesis is via inhibiting the Pol III repressor Maf1. These findings suggest that stimulation of tRNA synthesis may be one way that Ras promotes mRNA translation to drive cell and tissue growth.

## RESULTS

### Activation of Ras/ERK signalling leads to increased protein synthesis

We first examined whether Ras signalling regulates protein synthesis in *Drosophila* S2 cells using a puromycin-labelling assay. When a constitutively active Ras mutant (Ras^V12^) was expressed in *Drosophila* S2 cells using an inducible expression vector, we found an increase in protein synthesis, which was blocked by treatment of cells with cyclohexamide (CHX), an inhibitor of mRNA translation (Fig 1A). Also, using polysome profiling to measure mRNA translation, we saw an increase in polysome levels in Ras^V12^ overexpressing cells when compared with control cells (Fig 1B). Conversely when we blocked Ras/ERK signalling by treating cells with the MEK inhibitor, U0126, protein synthesis was decreased (Fig 1C). Finally, we found that total protein content/cell increased after Ras^V12^ was overexpressed in S2 cells (Fig 1D). Our findings suggest that one way that the Ras/ERK signalling pathway may drive growth in *Drosophila* is by promoting protein synthesis.

**Figure 1.**
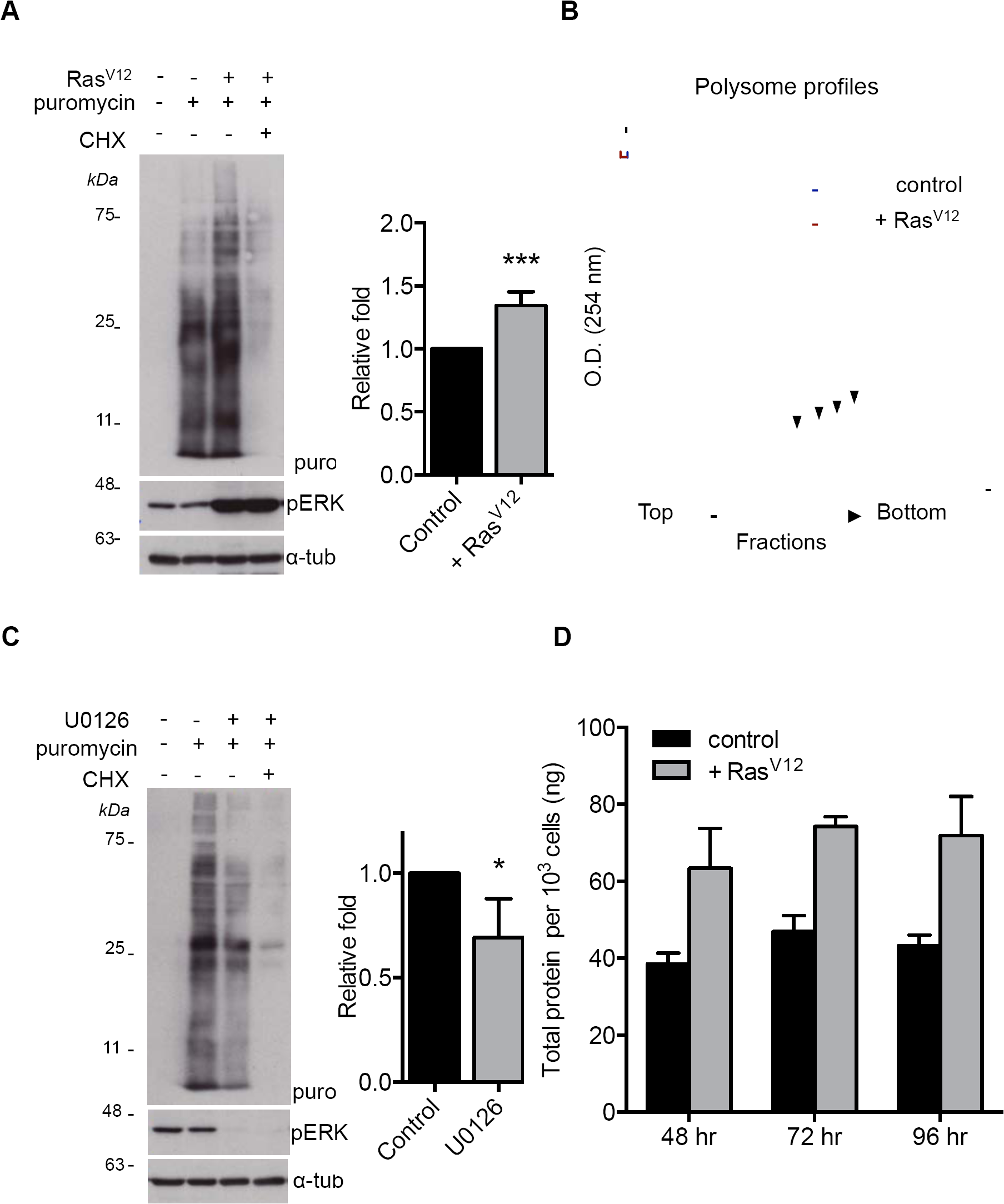
Activated Ras/ERK signalling pathway stimulates protein synthesis. (A) Ras^V12^ expression was induced in cultured *Drosophila* S2 cells for 24 hrs. Cells were then incubated in puromycin for 30 min. Protein extracts were separated by SDS-PAGE and analyzed by western blot with antibody to puromycin to measure the levels of puromycin-labelled peptides. Cyclohexamide treatment was for 15 mins prior to addition of puromycin. A phospho-ERK immunoblot is shown as an indication of Ras/ERK signalling pathway activation. An alpha-tubulin immunoblot is shown as a loading control (B) A Representative polysome profiles from control S2 cells (green) and S2 cells with induced Ras^V12^ expression (blue). Polysome peaks (arrowheads) in Ras^V12^ expressing cells were higher compared to controls, suggesting translation was increased. (C) *Drosophila* S2 cells were treated with in the presence or absence of 10 μM U0126 for 2 hours at 25°C. Cells were then incubated in puromycin for 30 min. Protein extracts were separated by SDS- PAGE and analyzed by western blot with antibody to puromycin to measure the levels of puromycin-labelled peptides. A phospho-ERK immunoblot is shown as an indication of Ras/ERK signalling pathway activation. An alpha-tubulin immunoblot is shown as a loading control (D) Total protein content per 10^3^ cells was calculated using a Bradford assay. All experiments were performed in at least three biological replicates and western blots were quantified using NIH Image J software. Quantifications are depicted in the bar graphs in A and B. Statistical comparisons were made with a Student’s t-test *** p < 0.0001, * p < 0.05

### Ras/ERK signalling promotes tRNA synthesis

We previously identified regulation of RNA Polymerase III and tRNA synthesis as a mechanism for controlling protein synthesis in *Drosophila* larvae (Marshall et al, 2012; Rideout et al, 2012). We showed that these processes were regulated by TORC1 kinase signalling, and that they were important for driving tissue and body growth(Marshall et al, 2012; Rideout et al, 2012). We were therefore interested in examining Ras signalling could also promote tRNA synthesis. We first used Northern blotting to examine tRNA levels in S2 cells. We found that Ras^V12^ overexpression lead to an increase in both pre-tRNA and mature tRNA levels, indicating enhanced tRNA synthesis (Fig 2A and Supplementary Fig S1A). In contrast, inhibiting Ras signalling with the MEK inhibitor U0126 blocked tRNA synthesis (Fig 2B and Supplementary Fig S1B). We saw similar results when we measured tRNA levels by qRT-PCR (Supplementary Fig S1C). We also examined the effects of Ras signalling on tRNA levels in the developing wing imaginal discs. Using *in situ* hybridization, we found that expression of *UAS-Ras*^*V12*^ in dorsal region of wing discs using the *apterous-GAL4* driver (*ap >Ras*^*V12*^) led to increased tRNA_i_^Met^ levels compared to cells in the ventral region. (Fig 2C). Similar results were obtained when we expressed either an activated form of Raf kinase (*UAS-Raf*^*gof*^) or *UAS-Ras*^*V12S35*^ (an effector loop mutant of Ras^V12^ that selectively activates the Raf-MEK-ERK branch of the Ras pathway) at the anterior-posterior boundary of wing discs using *dpp-GAL4* (Fig 2D).

**Figure 2.**
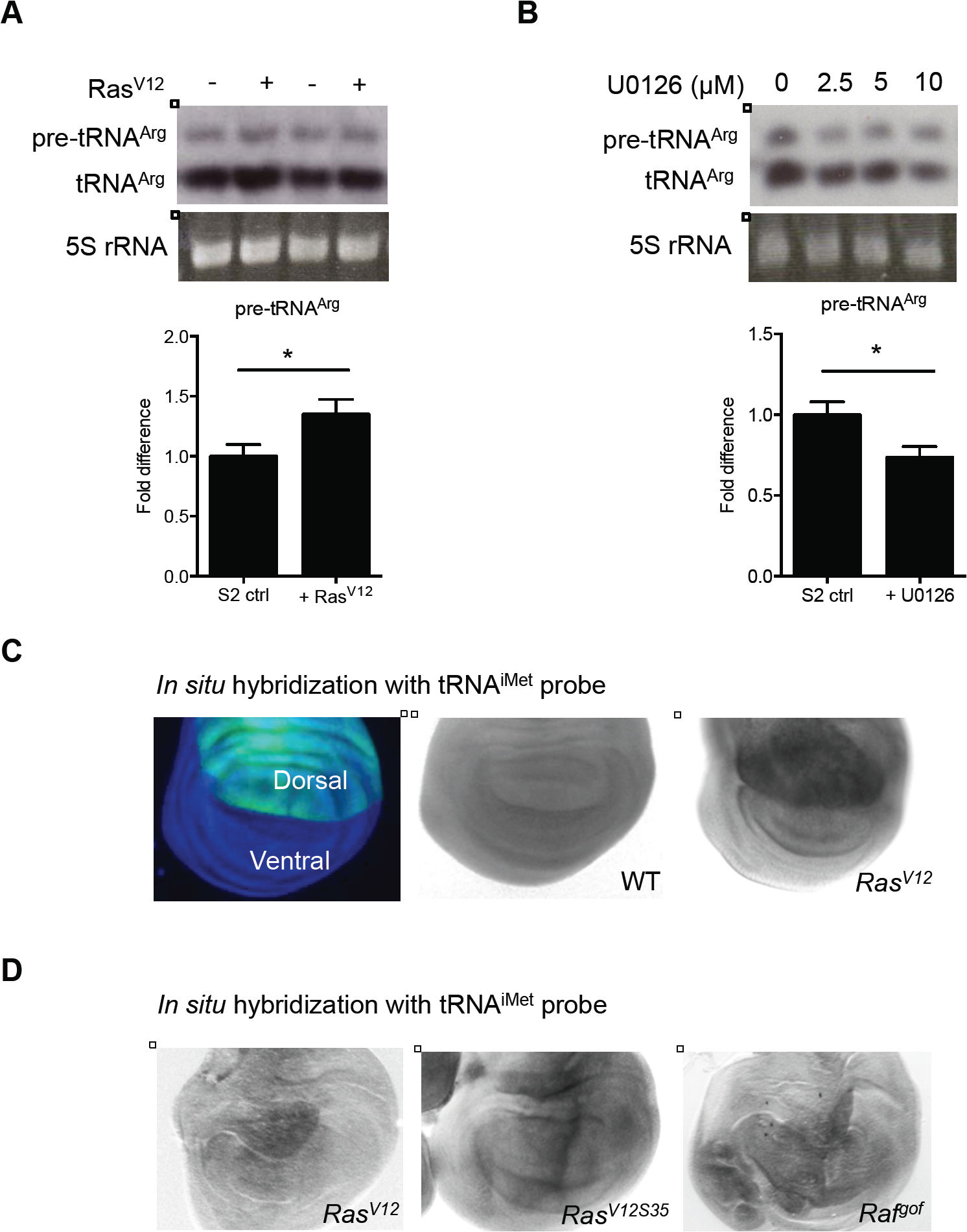
The Ras/ERK signalling pathway stimulates tRNA synthesis. (A) Ras^V12^ expression was induced in *Drosophila* S2 cells for 24 hours. Total RNA was isolated using Trizol reagent as described in the experimental procedures. 5 μg of total RNA per sample were separated on a 5% denaturing acrylamide gel and analyzed by Northern blotting. Levels of tRNA^Arg^ were detected using DIG-tRNA probes. Ethidium bromide stained 5S rRNA band was used as a loading control. Northern blots were quantified using NIH Image J software in the bar graphs. Student’s t-test, * p < 0.05. (B) *Drosophila* S2 cells were treated with 0, 2.5, 5 and 10 μM U0126 for 2 hours and then tRNA levels measured and quantified as in A. (C) An *ap-Ga¡4* driver was used to express *UAS-Ras*^*V12*^ in the dorsal region of larval wing imaginal discs. tRNA^iMet^ levels were detected by *in situ* hybridization using DIG-tRNA^iMet^ probe. Control discs were from flies carrying the *ap-GAL4* driver crossed to *w*^*1118*^ (D). A *dpp-Gal4* driver was used to express *UAS-EGFR, UAS-Ras*^*V12S35*^ or *UAS-Raf*^*gof*^ within a stripe of cells at the anterior-posterior border. tRNA^iMet^ levels were measured by in situ hybridization as in C.

### Brf1 is required for Ras-induced tRNA synthesis and cell proliferation in wing discs

Brf1 is a conserved component of TFIIIB complex, which is required for Pol III recruitment to tRNA genes (Geiduschek & Kassavetis, 2001). We previously showed that Brf1 is involved in controlling Pol III-dependent transcription, and tissue and body growth in *Drosophila* larvae (Marshall et al, 2012). Here we examined whether Brf1 is required for Ras-induced tRNA synthesis. We found that knocking down Brf1 using dsRNA in S2 cells (Supplementary Figure 2A, B) suppressed the Ras^V12^ induced increase in tRNA levels (Fig 3A). These data suggest that the elevation of tRNA levels upon Ras activation is due to increased Pol III transcription. We then examined whether Brf1 is required for Ras-induced growth in *Drosophila* wing discs. Overexpression of *UAS-EGFR* in the dorsal compartment of the wing imaginal disc (using an *apterous-Ga¡4* driver, *ap-GAL4*) stimulates Ras/ERK signalling and leads to tissue growth (Fig 3B and 3C). We found that RNAi-mediated knockdown of Brf1 by expression of a *UAS-Brf1* inverted repeat line (*UAS-Brf1 RNAi*) in the dorsal compartment had little effect on tissue growth. However, expression of *UAS-Brf1 RNAi* blocked the overgrowth seen with *UAS-EGFR* expression. Expression of *UAS-Brf1 RNAi* with ap-GAL4 had little effect on tissue growth, suggesting we are not knocking down Brf1 to a level that cannot support any growth. These data suggest that Brf1 is required for EGFR/Ras/ERK-mediated increases in epithelial tissue growth in *Drosophila*.

**Figure 3.**
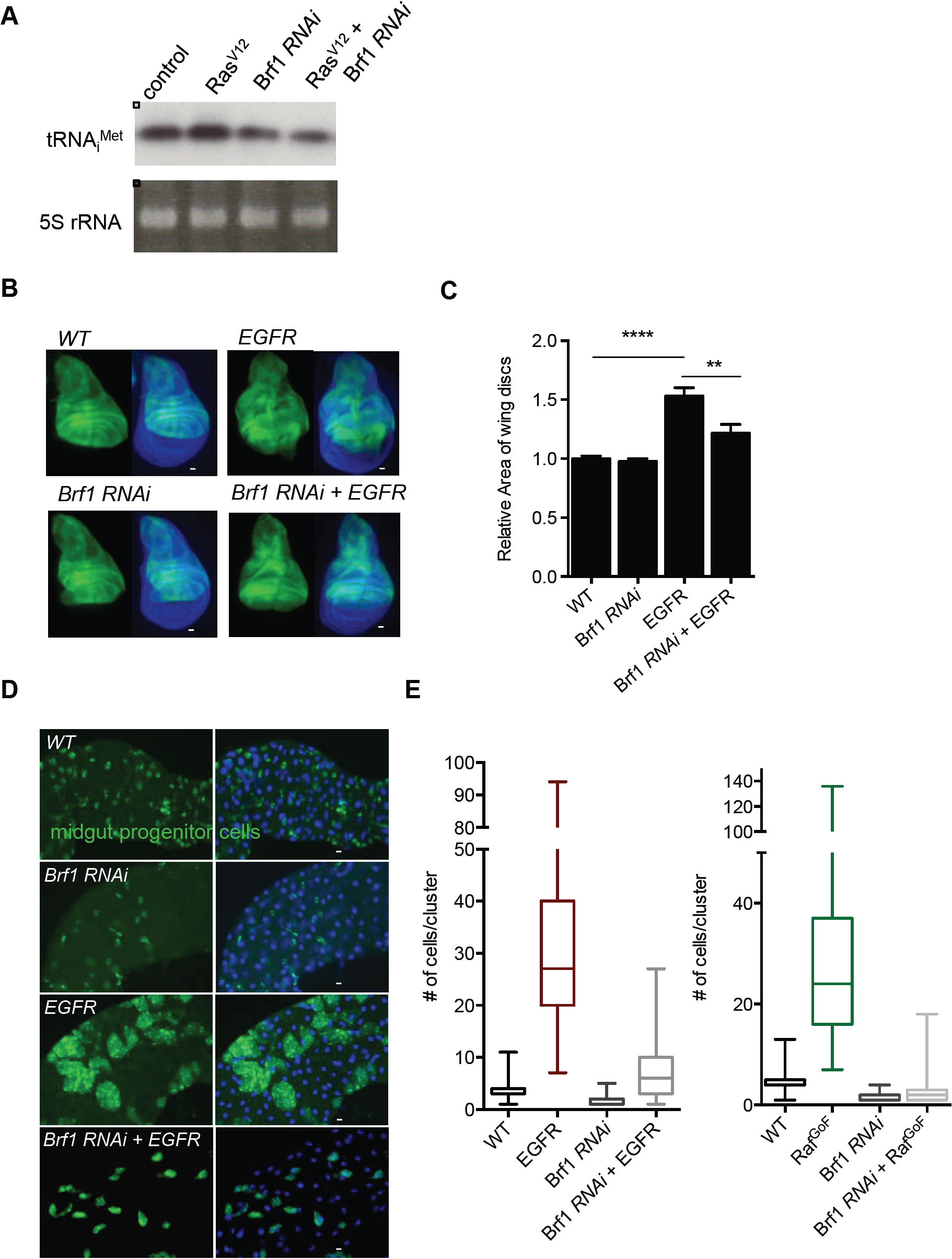
Brf1 is required for Ras-induced tRNA synthesis and growth in both wing imaginal discs and adult midgut progenitor cells (AMPs) (A). Ras^V12^ expression was induced in *Drosophila* S2 cells for 24 hours in either control cells or Brf1 knockdown cells, Brf1 was knocked down by incubating cells with dsRNA against Brf1. Control cells were treated with dsRNA to GFP. Total RNA was isolated with Trizol and analyzed by Northern blotting using a DIG-labelled antisense tRNA^iMet^ probe. Ethidium bromide stained 5S rRNA band was used as a loading control. (B, C) *UAS-EGFR* and *UAS-Brf1* were expressed, either alone or together, in the dorsal compartment of larval wing imaginal discs using an *ap-Ga¡4* driver. Control discs were from *ap-Ga¡4* crossed to w^111S^. Wing discs were dissected at the wandering L3 larval stage and the area of the GFP-marked dorsal compartment quantified using NIH imaging software (n > 50 wings per genotype, data presented as mean +/-SEM). Representative images are shown. (D) *UAS-EGFR* and *UAS-Brf1* were expressed, either alone or together, in the *Drosophila* larval AMPs using the *esg-Gal4*^*ts*^ system. Larvae were shifted to 29°C at 24 hrs of development to induce transgene expression and dissected as L3 larvae. AMPs are marked *by UAS-GFP* expression. DNA is stained with Hoechst dye (blue). (E) The number of cells in each AMP cluster was quantified for each of the genotypes in (D). Data are presented as box plots (25%, median and 75% values) with error bars indicating the min and max values.

### Brf1 is required for Ras/ERK-induced proliferation in adult mid-gut progenitor cells (AMPs) and adult intestinal stem cells (ISC)

A major role for the EGFR/Ras/ERK pathway is in the growth and maintenance of the *Drosophila* intestine. In larvae, activation of the pathway plays a central role in controlling the proliferation of adult midgut progenitor cells (AMPs), which eventually give rise to the adult intestine (Jiang & Edgar, 2009). In the adult the EGFR/Ras/ERK pathway is required to promote stem cell proliferation and tissue regeneration (Biteau & Jasper, 2011; Buchon et al, 2010; Cordero et al, 2012; Jiang et al, 2011; Xu et al, 2011). We therefore examined whether Brf1-mediated Pol III transcription was required for these proliferative effects of Ras/ERK signalling. We first examined the larval intestine. During the larval period, AMPs proliferate and give rise to clusters of ~5-10 cells scattered throughout the larval intestine. These cell clusters eventually proliferate and fuse during metamorphosis to give rise to the adult intestinal epithelium. The EGFR/Ras/ERK pathway controls the proliferation of AMPS. Overexpression of either *UAS-EGFR* or *UAS-Raf*^*gof*^ in the AMPs using the temperature-sensitive *escargot-Gal4 (esg-Gal4*^*ts*^) system lead to a massive increase AMP proliferation and an increase in the numbers of AMP cells per cluster as previously reported. We found that expression of *UAS-Brf1 RNAi* (Fig 3D and 3E and Supplementary Fig S2C) lead to a small reduction in the number of cells per cluster. However, we found that when co-expressed along with *UAS-EGFR* or *UAS-Raf*^*gof*^*, UAS-Brf1 RNAi* blocked the increase in AMP cell number seen. These data indicate Brf1 is required for EGFR/Ras/ERK mediated cell proliferation.

We next examined Brf1 function in homeostatic growth in the adult intestine. Damage to intestinal epithelial cells leads to an increase in expression and release of EGF ligands from both intestinal cells and underlying visceral muscle. These EGF ligands then act on the intestinal stem cells (ISCs) to stimulate the Ras/ERK pathway, which triggers stem cell growth and division, and promotes regeneration of the intestinal epithelium. This damage-induced increase in ISC proliferation is dependent on EGFR/Ras/ERK signalling and can be mimicked by genetically activating the pathway specifically in the stem cells. We tested a requirement for Brf1 in this Ras-mediated homeostatic growth response. We first examined the effects of intestinal damage. As previously reported (Biteau & Jasper, 2011), we found that feeding flies either DSS or bleomycin - two different gut stressors - leads to an increase in ISC proliferation. However, we found that this effect was inhibited when we knocked down Brf1 (using *UAS-Brf1 RNAi* expression) specifically in the ISCs and their transient daughter cells, the enteroblasts (EBs) using the inducible *esg-Gal4*^*ts*^ system (Fig 4A and 4B). We next examined the effects of activation of the Ras/ERK pathway. As previously reported, when we overactivated the pathway in stem cells by expressing *UAS-Raf*^*gof*^ using *esg-Gal4*^*ts*^, we saw an increase cell proliferation as indicated by a marked increased in GFP labelled ISCs and EBs (Fig 4C). Expression of a *UAS-Brf1 RNAi* had little effect on GFP labelled cells, but when co-expressed with *UAS-Raf*^*gof*^ it blocked the increase in cell proliferation. These results suggest that Brf1 and Pol III-dependent transcription is required for stem cell proliferation in the adult intestine.

**Figure 4.**
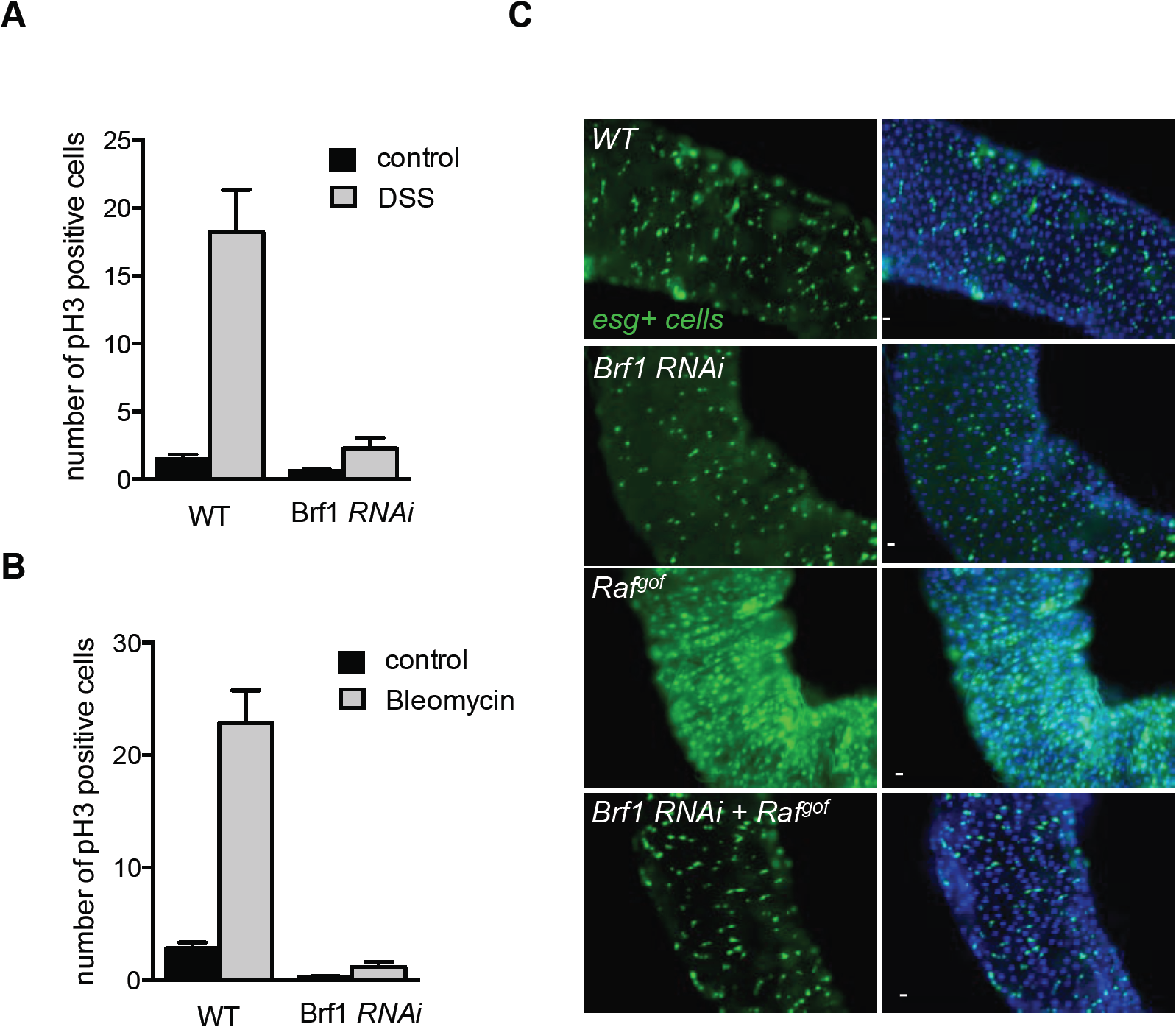
Brf1 is required for in intestinal stem cells (ISCs) homeostasis and for Ras-induced cell proliferation. (A, B) UAS-Brf1 RNAi was expressed in adult ISCs and EBs using the *esg-GAL4*^*ts*^ system. Control flies were *esg-GAL4*^*ts*^ flies crossed to w^111S^. Flies were then fed with sucrose or sucrose plus DSS (A) or Bleomycin (B) for 2 days. Intestines were then dissected and stained for phospho-histone H3 positive cells. Data present mean number of phospho-histone H3 cells per intestine +/ SEM. N >15 intestines per condition. (C) *UAS-Raf*^*gof*^ and *UAS-Brf1* were expressed, either alone or together, in the adult ISCs and EBs using the *esg-Gal4*^*ts*^ system. Esg positive cells are marked with GFP and DNA is stained with Hoechst dye. Knockdown of Brf *1(UAS-Brf RNAi*) suppresses the increased proliferation seen with *UAS-Raf*^*gof*^ expression.

### dMyc is required but not sufficient for Ras-induced tRNA synthesis

We next wanted to examine how Ras signalling stimulates Pol III-dependent tRNA transcription. One candidate regulator we tested was dMyc. In both mammalian cells and *Drosophila*, dMyc can interact with Brf1 and stimulate Pol III-dependent transcription (Gomez-Roman et al, 2003; Marshall et al, 2012; Steiger et al, 2008). Moreover, studies in both mammalian cells and *Drosophila* suggest Ras signalling can regulate Myc levels and that Myc is required for Ras-induced growth (Prober & Edgar, 2000; Prober & Edgar, 2002; Ren et al, 2013; Sears et al, 1999; Soucek et al, 2013). Indeed, we found that the *UAS-EGFR-* and *UAS-Ras*^*v12S55*^-induced proliferation of larval AMPs was blocked when we knocked down dMyc by expression of a *UAS-dMyc RNAi* construct (Supplementary Fig S3A-C). We therefore examined whether dMyc functions downstream of Ras in the control of Pol III. Using S2 cells we found that the increase in tRNA levels seen following Ras^V12^ expression was blocked when cells were treated with dsRNA to knockdown dMyc (Fig 5A). In contrast we found that overexpression of dMyc in S2 cells was not able to induce tRNA synthesis in cells in which the Ras pathway was inhibited by treatment with the MEK inhibitor UO126 (Fig 5B). Together, these data indicate that dMyc is required, but not sufficient, to mediate the effects of Ras signalling on tRNA synthesis.

**Figure 5.**
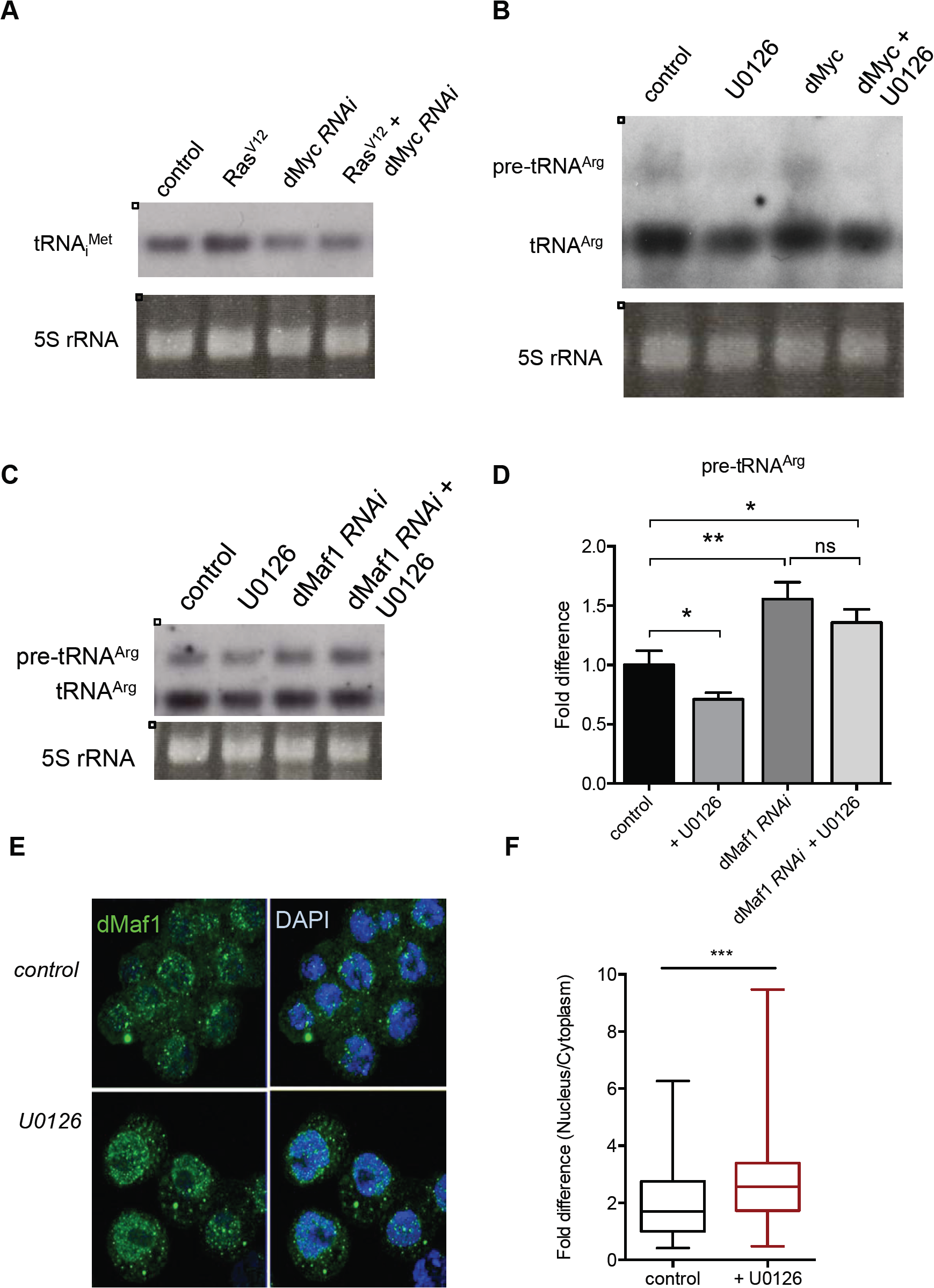
Ras induces tRNA synthesis by via inhibition of the Pol III repressor, Mafl. (A) Ras^V12^ expression was induced in *Drosophila* S2 cells for 24 hours in either control cells or dMyc knockdown cells. dMyc was knocked down by incubating cells with dsRNA against dMyc. Control cells were treated with dsRNA to GFP. Total RNA was isolated with Trizol and analyzed by Northern blotting using a DIG-labelled antisense tRNA^iMet^ probe. Ethidium bromide stained 5S rRNA band was used as a loading control. (B) dMyc expression was induced in S2 cells for 24hrs, and then cells were treated with 10 μM U0126 or DMSO for 2 hours. Total RNA was isolated with Trizol and analyzed by Northern blotting using DIG-labelled tRNA^Arg^ probe. Ethidium bromide stained 5S rRNA band was used as a loading control. (C) dMaf1 was knocked down in *Drosophila* S2 cells by incubating cells with dsRNAs against *dMafl.* Control cells were treated with dsRNA to GFP. Cells were then treated with DMSO (control) or 10 μM U0126 for 2 hrs. Total RNA was isolated with Trizol and analyzed by northern blot using DIG-labelled tRNA^Arg^ probe. Ethidium bromide stained 5S rRNA band was used as a loading control. (D) Northern blots were quantified using NIH Image J software. The bar graphs represent the mean fold difference (+/-SEM) in pre-tRNA band intensity compared to control group. Statistical differences between each group the control group were assessed using Student’s t-test, **p<0.01, * p < 0.05. (E) dMaf1 subcellular localization was assessed by immunostaining with an anti-dMaf1 antibody in both control and 10 μM U0126 treated S2 cells. Green: dMaf1 staining; blue: Hoechst-stained nuclei. (E) Differential localization of dMaf1 was quantified by measuring the area of the cytoplasm and nucleus in the confocal sections. Ratio of nucleus/cytoplasm is quantified using NIH Image J and plotted in the graph as fold difference vs. control. Data are presented as box plots (25%, median and 75% values) with error bars indicating the min and max values. Statistical differences were assessed using a Student’s t-test, *** p<0.001.

### Ras signalling promotes tRNA synthesis by inhibiting the RNA poi III repressor dMafl

Another candidate that we considered as a mediator of Ras-induced tRNA synthesis is the conserved Pol III repressor, Maf1. Studies in yeast, *Drosophila* and mammalian cells have shown that inhibition of Maf1 is the main way that the nutrient-dependent TORC1 kinase pathway stimulates Pol III and tRNA synthesis (Kantidakis et al, 2010; Marshall et al, 2012; Michels et al, 2010; Shor et al, 2010; Wei et al, 2009). Knockdown of dMaf1 has been shown to promote tRNA synthesis, and to enhance tissue and body growth in *Drosophila* (Rideout et al, 2012). Here, we found that when we expressed *UAS-dMaf1 RNAi* in the Ras-responsive AMP cells during larval development using *esg-GAL4*^*ts*^, we observed a modest, but significant increase in the number of AMP cells per cluster (Supplementary Fig S4A). Although considerably weaker than the effect of Ras pathway activation, this effect of dMaf1 knockdown was similar to the increase in AMP numbers seen with overexpression of dMyc, another stimulator of tRNA synthesis and mRNA translation (Supplementary Fig S4B). We therefore next examined whether the Ras/ERK pathway functions to promote tRNA synthesis by inhibiting dMaf1. We examined tRNA levels by Northern blot in S2 cells, and, as described above, we saw that treatment of cells with the MEK inhibitor UO126 led to reduced tRNA synthesis (Fig 5C and 5D and Supplementary Fig S4). However, we found that this decrease in tRNA synthesis was reversed when cells were treated with dsRNA to knockdown dMaf1 levels (Fig 5C and 5D and Supplementary Fig S4). These data suggest that a main way that Ras/Erk signalling functions to promote tRNA synthesis is by inhibiting the Pol III repressor function of dMaf1. Studies in both yeast and mammals suggest that one way that dMaf1 can be regulated is by controlling it’s nuclear localization. We tested this in S2 cells using an antibody to endogenous dMaf1. Under our normal media culture conditions, we observed that dMaf1 was localized throughout the cell (Fig 5E and 5F). However, treatment of cells with the MEK inhibitor U0126 leads to a significant increase in nuclear localization of dMaf1. Thus, Ras/ERK signalling functions to prevent nuclear accumulation of dMaf1, hence blocking its Pol III repressor activity and promoting tRNA synthesis.

## DISCUSSION

We propose that stimulation of RNA polymerase III and tRNA synthesis contributes to the ability of the conserved Ras/ERK pathway to promotes mRNA translation and growth. Our data indicate that the main way Ras controls Pol III is by inhibiting the Maf1 repressor, in large part by preventing its nuclear accumulation. Maf1 is a phospho protein and studies in yeast and mammalian cells have described how phosphorylation can regulate Maf1 nuclear localization. For example, both TORC1 and PKA can phosphorylate on several conserved residues(Huber et al, 2009; Kantidakis et al, 2010; Michels et al, 2010; Moir et al, 2006; Shor et al, 2010; Wei et al, 2009). This phosphorylation prevents Maf1 nuclear accumulation and allows both kinases to stimulate Pol III. In contrast, dephosphorylation of Maf1 by both PP2A and PP4 protein phosphatases leads to nuclear accumulation of Maf1 and Pol III repression (Oficjalska-Pham et al, 2006; Oler & Cairns, 2012; Roberts et al, 2006). Thus it is possible that ERK may function by promoting Maf1 phosphorylation - either directly or indirectly - to prevent its function. Other mechanisms may also be important for Ras to simulate tRNA synthesis. For example, one study in mammalian cells showed that ERK could phosphorylate and regulate Brf1 function (Felton-Edkins et al, 2003). Also Ras was shown to upregulate TBP, which can increase transcription by all three RNA polymerases(Zhong et al, 2004). Importantly, Maf1 function is conserved suggesting that Ras/ERK-dependent regulation of Maf1 and tRNA synthesis that we describe in *Drosophila* may operate in other organisms, particularly human cells.

We also show that the transcription factor Myc is required for the effects of Ras on Pol III and tRNA synthesis. Previous work from both mammalian cells and *Drosophila* has shown that in some cells Ras can promote Myc levels and that Ras-mediated growth requires Myc function(Prober & Edgar, 2000; Prober & Edgar, 2002; Ren et al, 2013; Sears et al, 1999; Soucek et al, 2013). We previously showed that *Drosophila* Myc could stimulate expression of the Pol III transcription factor, Brf1, and also other Pol III subunits (Marshall et al, 2012). In addition, Myc can directly interact with Brf1 and localize at Pol III to directly stimulate tRNA transcription (Gomez-Roman et al, 2003; Marshall et al, 2012; Steiger et al, 2008). We suggest that both these effects are under the upstream control of Ras/ERK signalling and may, in part, explain the requirements for Myc in Ras-induced growth in both animal development and cancer.

Previous studies in mammalian cells have shown that Ras/ERK signalling can promote protein synthesis by stimulating translation initiation factor function. We suggest that inhibition of Maf1 represents another target of Ras/ERK signalling, and that the subsequent increase in tRNA levels may cooperate with enhanced translation initiation factor activity to promote maximal stimulation of mRNA translation. Most of the work on Ras-mediated gene expression has focused on the effect of several Pol II transcription factors identified downstream of Ras in *Drosophila* such as *fos, pointed*, and *capicua*(Baonza et al, 2002; Biteau & Jasper, 2011; Jin et al, 2015; Tseng et al, 2007). Stimulation of Pol III transcription to enhance tRNA levels and mRNA translation may provide another layer of control on overall gene expression by Ras signalling. For example translational control of cell cycle genes has been proposed as one way to couple growth signalling pathways to cellular proliferation (Dowling et al, 2010; Polymenis & Schmidt, 1997). Furthermore, selective translational regulation of certain mRNAs has been shown to regulate growth and metastatic behaviour of tumour cells (Hsieh et al, 2012; Morita et al, 2013; Thoreen et al, 2012).

Ras is one of the most often overactivated or mutated pathways in cancer, hence our findings may also have implications for processes that contribute to tumour growth and metastasis. Indeed, there is increasing appreciation for potential roles for alterations in tRNA biology in cancer cells (Grewal, 2015). For example, tRNA expression profiling has revealed that levels of many tRNAs are elevated in different cancer types (Pavon-Eternod et al, 2009; Zhou et al, 2009). Interestingly, these changes in tRNA levels have been shown to correlate with codon usage in mRNAs whose expression also changes in cancer cells (Gingold et al, 2014). Several studies have reported that increasing the levels of specific tRNAs can promote tumour growth and metastatic behavior (Birch et al, 2016; Clarke et al, 2016; Goodarzi et al, 2016; Pavon-Eternod et al, 2013). Previous work also showed that increasing tRNA levels alone is sufficient to drive growth in *Drosophila* (Rideout et al, 2012; Rojas-Benitez et al, 2015). Hence, an increase in tRNA levels caused by oncogenic Ras signalling may be a driver of tumour growth and progression, rather than simply a consequence of increased growth. Ras also controls other process such as cell fate specification, differentiation and cell survival. Many of these effects are mediated through translation and so may also rely on the effects of Ras on tRNA synthesis.

## EXPERIMENTAL PROCEDURE

### *Drosophila* stocks

Flies were raised on standard medium (150 g agar, 1600 g cornmeal, 770 g Torula yeast, 675 g sucrose, 2340 g D-glucose, 240 ml acid mixture (propionic acid/phosphoric acid) per 34 L water) and maintained at 25°C, unless otherwise indicated. The following fly stocks were used:

*w*^*1118*^,

*yw*,

*UAS-Ras*^*V12*^*, (Prober & Edgar, 2002*)

*UAS-Ras*^*V12*^*S35, (Prober & Edgar, 2002*)

*UAS-EGFR, (Prober & Edgar, 2002*)

*UAS-Rafi*^*of*^*, (Prober & Edgar, 2002*)

*UAS-Brf RNAi (NIG, Japan*),

*UAS-dMyc (Grewal et al, 2005*)

*UAS-Maf1 RNAi*,

*esg-gal4, tub-GAL80ts, UAS-GFP, (Jiang et al, 2011*)

*ap-gal4*,

*dpp-gal4*.

For all GAL4/UAS experiments, homozygous GAL4 lines were crossed to the relevant UAS line(s) and the larval or adult progeny were analyzed. Control animals were obtained by crossing the rrelevant homozygous GAL4 line to either *w*^*1118*^ or *yw* depending on the genetic background of the particular experimental UAS transgene line.

### Cell Culture and Transfection

*Drosophila* Schneider S2 cells were grown at 25°C in Schneider’s medium (Gibco; 11720-034) supplemented with 10% fetal bovine serum (Gibco; 10082-139), 100 U/ml penicillin and 100 U/ml streptomycin (Gibco; 15140). Stably transfected inducible Ras^V12^ cells were a gift from the lab of Marc Therrien (Ashton-Beaucage et al, 2014). Stably transfected inducible dMyc cells were a gift from the lab of Paula Bellosta (Bellosta et al, 2005). Both Ras^V12^ and dMyc expression are under the control of a metallothionein promoter. For all experiments Ras^V12^ or dMyc were induced by addition of copper sulphate to the culture media.

*dsRNA Treatment of S2 cells:*. Cells were pretreated with 15 μg of dsRNAs in the absence of 1 ml of serum for 30 mins and then 2 mls of serum was added and incubated for 96 to 120 hrs. Control cells were treated with ds RNA to Green Fluorescent Protein (GFP). Cells were harvested by centrifugation at 4 °C and washed with cold PBS and frozen for RNA isolation or protein extraction. *MEK inhibitor (U0126) treatment of Drosophila S2 cells:* S2 cells were cultured at 25°C in Schneider’s medium (Gibco; 11720-034) supplemented with 10% fetal bovine serum (Gibco; 10082-139), 100 U/ml penicillin and 100 U/ml streptomycin (Gibco; 15140). Cells were treated with either 10 μΜ U0126 (Promega Cat. No. V1121) or DMSO (Sigma; D2650) for 2 hours. Then cells were washed twice with ice-cold PBS. Cells were then used to isolate RNA or make protein extracts as described below.

### Preparation of protein extracts

*Drosophila* S2 cells were lysed with a buffer containing 20 mM Tris-HCl (pH 8.0), 137 mM NaCl, 1 mM EDTA, 25 % glycerol, 1% NP-40 and with following inhibitors 50 mM NaF, 1 mM PMSF, 1 mM DTT, 5 mM sodium ortho vanadate (Na3VO4) and Protease Inhibitor cocktail (Roche Cat. No. 04693124001) and Phosphatase inhibitor (Roche Cat. No. 04906845001) according to the manufacturer’s instruction.

### Western Blot and Antibodies

Protein concentrations were measured using the Bio-Rad Dc Protein Assay kit II (5000112). Protein lysates (15 μg to 30μl) were resolved by SDS-PAGE and electrotransferred to a nitrocellulose membrane, subjected to Western blot analysis with specific antibodies, and visualized by chemiluminescence (enhanced ECL solution (Perkin Elmer). Brf primary antibodies were against a C-terminal fragment of *Drosophila* Brf, alpha-tubulin (E7, *Drosophila* Studies Hybridoma Bank), dMyc(Prober & Edgar, 2002), phospho-ERK (Cell Signalling Technology 4370) and ERK(Cell Signalling Technology 4695). Peptide antiserum against *Drosophila* Maf1 was raised by immunizing rabbits with synthetic peptide LADFSPNFRC corresponding to residues 65-74 (GL Biochem (Shanghai) Ltd).

### Puromycin-labelling protein synthesis assay

10 μM puromycin was added to *Drosophila* S2 cell culture media and the cells were incubated with puromycin for 30 min at 25 °C. Cells were harvested by centrifugation at 4°C and washed with cold PBS. Cells were frozen on dry ice and then lysed according to the Western blot protocol described above and analyzed by SDS-PAGE and western blotting using an anti-puromycin antibody (3RH11) (Kerafast, Catalog No.EQ0001) at 1:2000 dilution.

### In situ hybridization in wing imaginal discs

*In situ* hybridization analysis of tRNA^iMet^ in wing discs was performed using digoxigenin-labelled probes. The digoxigenin-labelled antisense riboprobe for tRNA^iMet^ was synthesized by *in vitro* transcription using Roche digoxigenin labeling kit. The tRNA^iMet^ was amplified from either genomic DNA or an expression plasmid containing the tRNA^iMet^ gene. Primers used for the amplification are listed in Supplementary Table S1. The in situ hybridizations were performed as described previously(Grewal et al, 2007). Briefly, inverted larvae were fixed in 4% paraformaldehyde (PFA), permeablized and then hybridized with probe overnight at 55 °C (hybridization buffer contained 50% formamide, 5xSSC pH 6.2, 100 μg/ml salmon sperm DNA, 50 μg/ml heparin and 0.1% Tween 20). Probe binding was detected using anti-DIG-AP fab fragments (1:2000) in 1x PBTw and staining with using 2% 4-nitroblue tetrazolium chloride (NBT), and 5-bromo-4-chloro-3-indolyl-phosphate (BCIP) detection system (Roche).

### Northern blot analysis

Total RNA was extracted from *Drosophila* S2 cells using TRIzol. 5 μg total RNA was separated on a 5 % denaturing polyacrylamide/urea gel and northern blotting was carried using alkaline transfer. Hybridization of tRNA probes were carried out as described in Roche DIG Easy Hyb (Cat. No.11603558001). Digoxigenin-labelled probes were made by *in vitro* transcription using either full-length cDNAs or PCR fragments as templates.

### Immunostaining

*Drosophila* S2 cells were fixed in 4% paraformaldehyde at room temperature for 20 mins on cover slips. Wash with 1x PBS and permeabilized with 0.1% Triton X in PBS by washing 2x for 5 mins. Cells were blocked with 5% FBS, 0.1% Triton X in PBS for 2 hours. Primary Maf1 antibody was diluted in 5% BSA in PBS at 1:500 dilution and incubated overnight at 4°C. Then washed 3x with 0.1% Triton X for 5 min each and Alexa 568 (Molecular probes) goat-anti rabbit secondary antibody was diluted at 1:400 in 5% BSA in PBS for 2 hours at room temperature. Then, cells were washed 3x with 0.1 % Triton X in PBS for 5 min each and mounted using Vector Shield mounting medium.

### Real-Time Quantitative PCR

Total RNA was extracted using TRIzol according to manufacturer’s instructions (Invitrogen; 15596018). RNA samples were DNase treated according to manufacturer’s instructions (Ambion; 2238G) and reverse transcribed using Superscript II (Invitrogen; 100004925). The generated cDNA was used as a template to perform qRT-PCRs (ABI 7500 real time PCR system using SyBr Green PCR mix) using specific primer pairs (sequences available upon request). PCR data were normalized to the average fold change of either actin or Glyceraldehyde-3-phosphate dehydrogenase (GAPDH). Each experiment was independently repeated a minimum of three times. All data were analysed by Student’s t-tests.

### Polysome gradient centrifugation

Polysome gradient centrifugation was performed as described (Rideout et al, 2012). 100 million *Drosophila* S2 cells were lysed in 1 ml of lysis buffer (25 mM Tris pH 7.4, 10 mM MgCl2, 250 mM NaCl, 1% Triton X-100, 0.5% sodium deoxycholate, 0.5 mM DTT, 100 mg/ml cycloheximide, 1 mg/ml heparin, Complete mini Roche protease inhibitor (Roche), 2.5 mM PMSF, 5 mM sodium fluoride, 1 mM sodium orthovanadate and 200 U/ml ribolock RNAse inhibitor (Fermentas) using a Dounce homogenizer. The lysates were centrifuged at 15,000 rpm for 20 minutes and the supernatant was removed carefully. 150 to 250 g μg RNA was layered gently on top of a 15–45% w/w sucrose gradient (made using 25 mM Tris pH 7.4, 10 mM MgCl2, 250 mM NaCl, 1 mg/ml heparin, 100 mg/ml cycloheximide in 12 ml polyallomer tube) and centrifuged at 37,000 rpm for 150 minutes in a Beckmann Coulter Optima L-90K ultracentrifuge using a SW-41 rotor. Polysome profiles were obtained by pushing the gradient using 70% w/v Sucrose pumped at 1.5 ml/min into a continuous OD254 nm reader (ISCO UA6 UV detector) showing the OD corresponding to the RNA present from the top to the bottom of the gradient.

## Author contributions

S. S-P. and S.S.G. designed research; S.S-P., B.L., R.J. and S.S.G. performed experiments, analyzed the data and S. S-P and S.S.G wrote the paper.

## Acknowledgements

We thank Dr. Marc Therrien and Dr. Paola Bellosta for *Drosophila* S2 cells stably transfected with Ras^V12^ and Myc cell lines. We thank Bloomington Stock Centres for providing flies, Dr. Shinako Takada for providing the Brf antibody, Dr. Paola Bellosta for the Myc antibody. This work was supported by a Canadian Institutes of Health Research (CIHR) grant to S.S.G. S S-P was supported by a postdoctoral fellowship from Alberta Innovates Health Solutions (AIHS).

---

**Supplementary Figure S1.**
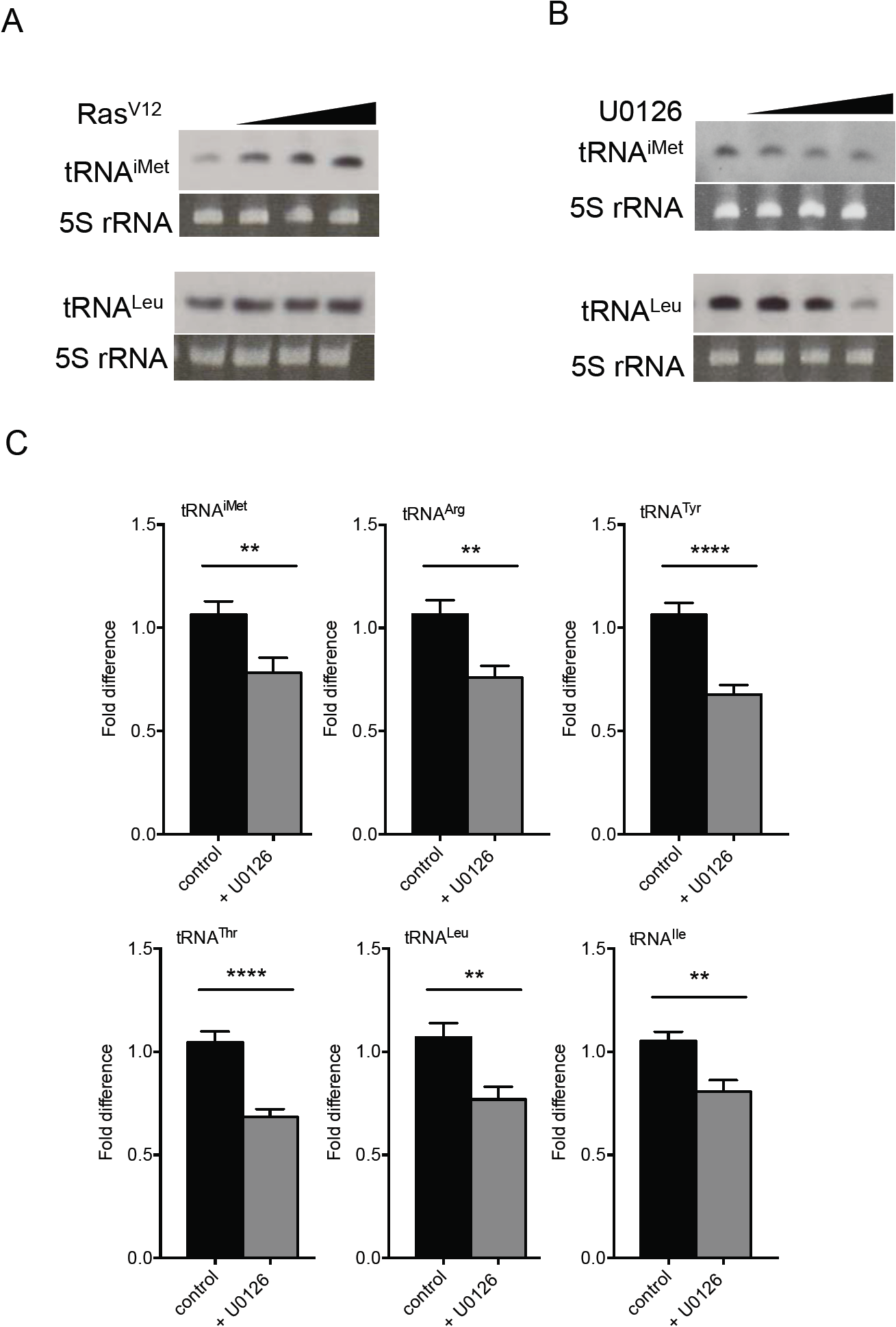
Ras/ERK signalling pathway regulates tRNA synthesis(related to Figure 2) (A and B) Ras^V12^ expression was induced in Drosophila S2 cells. Total RNA was isolated and analyzed by Northern blot. Levels of tRNA^iMet^ and tRNA^Leu^ were detected using DIG-labelled tRNA probes. Eithidium bromide stained 5S rRNA band was used as a loading control. (B) S2 cells were treated with U0126, and total RNA was isolated and analyzed by qRT-PCR.

**Supplementary Figure S2.**
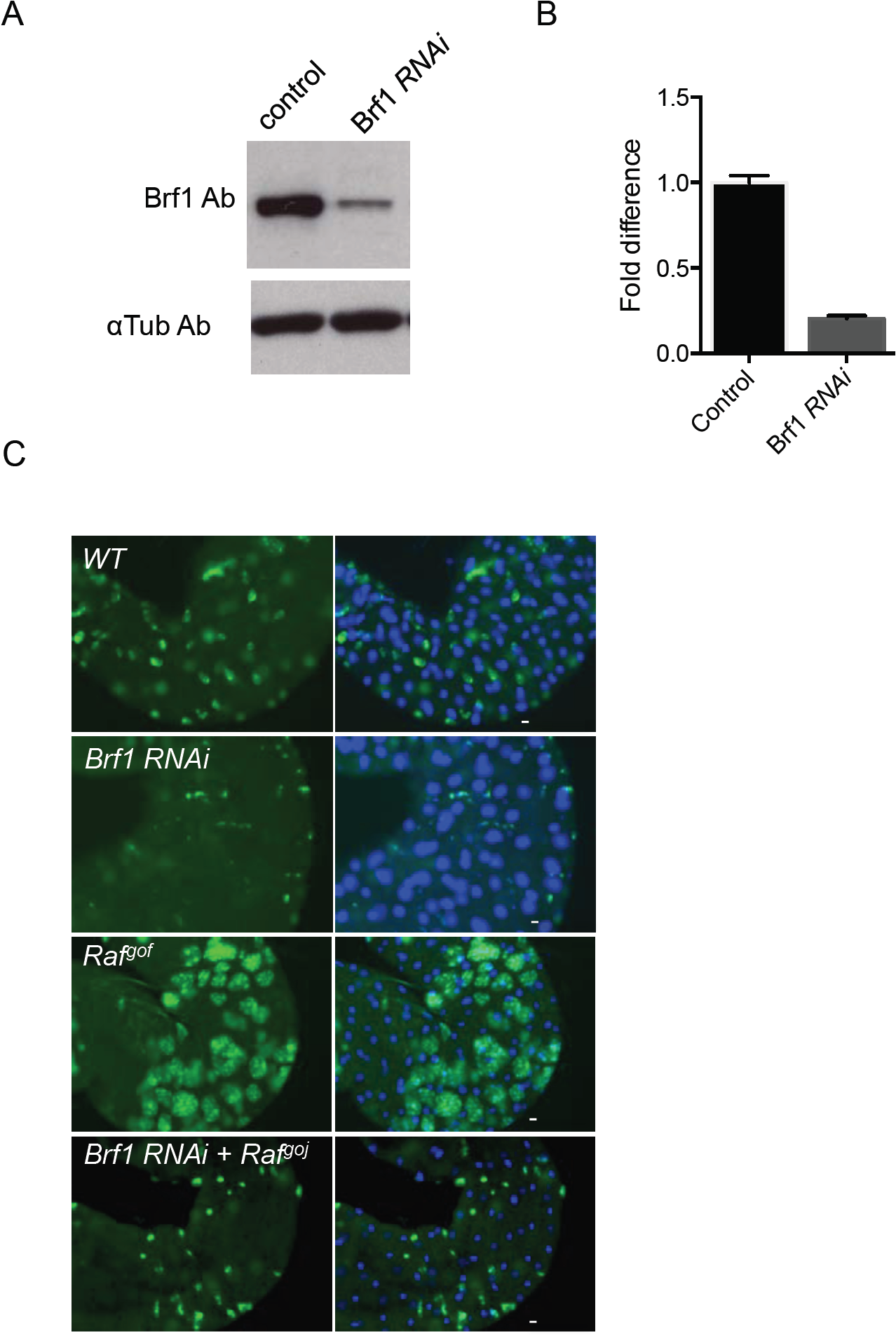
Brf1 is required for Ras-induced cell proliferation in AMPs(related to Figure 3) (A) Brf1 protein levels were measured by Western blot in Drosophila S2 cells treated with dsRNA against Brf1. Control cells were treated with GFP dsRNA (B) Brf1 mRNA levels were measured by qRT-PCR in Drosophila S2 cells treated with dsRNA against Brf1 or GFP (control). (C) *UAS-RaJ*^*gof*^ and *UAS-Brf1* were expressed, either alone or together, in the Drosophila larval AMPs using the *esg-Gal4*^*ts*^ system. Larvae were shifted to 29°C at 24 hrs of development to induce transgene expression and dissected as L3 larvae. AMPs are marked *by UAS-GFP* expression. DNA is stained with Hoechst dye (blue) Representative images are shown for each genotype.

**Supplementary Figure S3.**
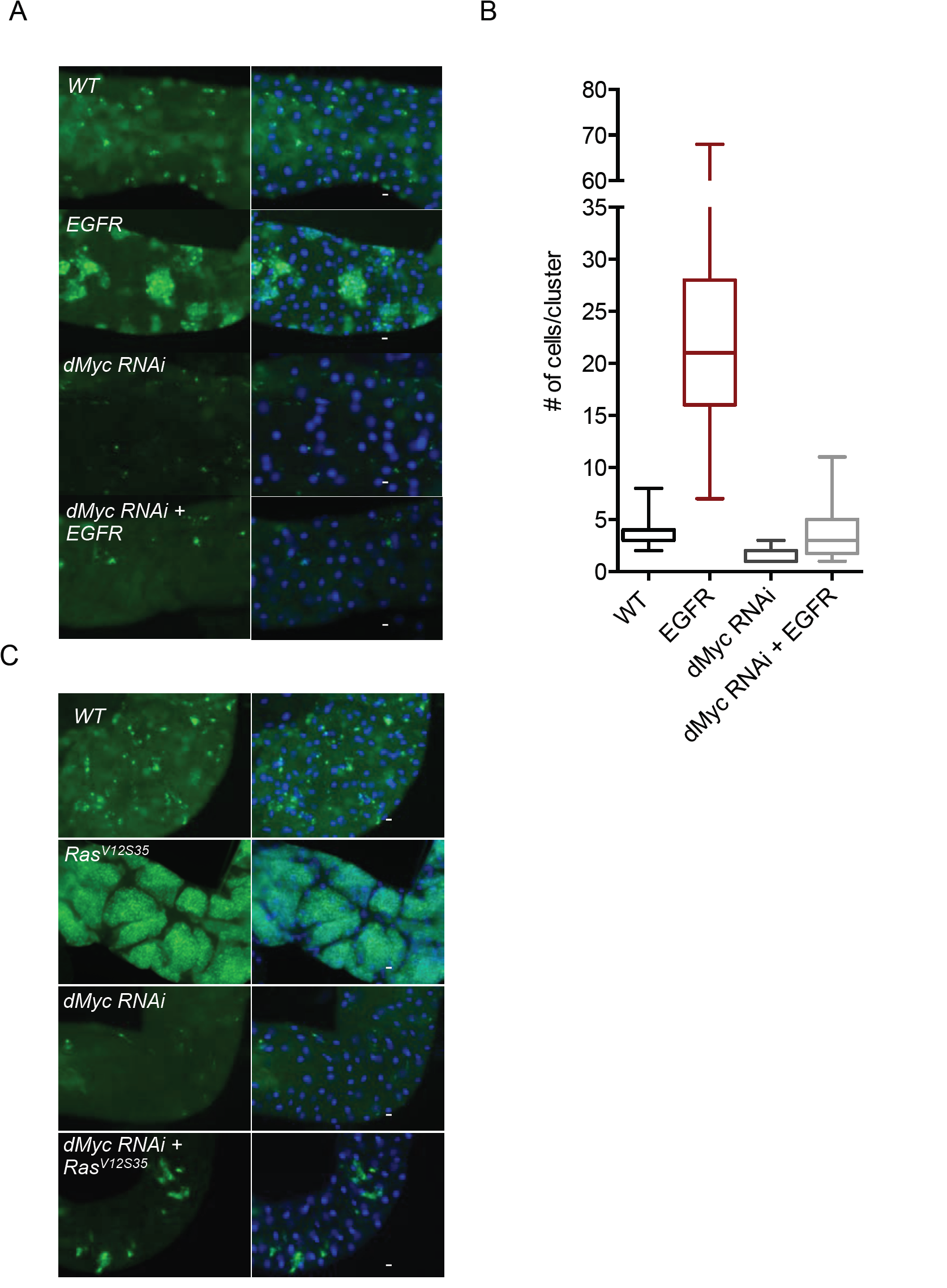
dMyc is required for Ras-induced AMP cell proliferation(related to Figure 5) *UAS-EGFR* (A) or *UAS-Ras*^*V12S35*^ were expressed, either alone or together with *UAS-dMyc RNAi* in the Drosophila larval AMPs using the *esg-Gal4*^*ts*^ system. Larvae were shifted to 29°C at 24 hrs of development to induce transgene expression and dissected as L3 larvae. AMPs are marked by *UAS-GFP* expression. DNA is stained with Hoechst dye (blue) (B) Numbers of cells in each AMP cluster were counted and expressed as box plots.

**Supplementary Figure S4.**
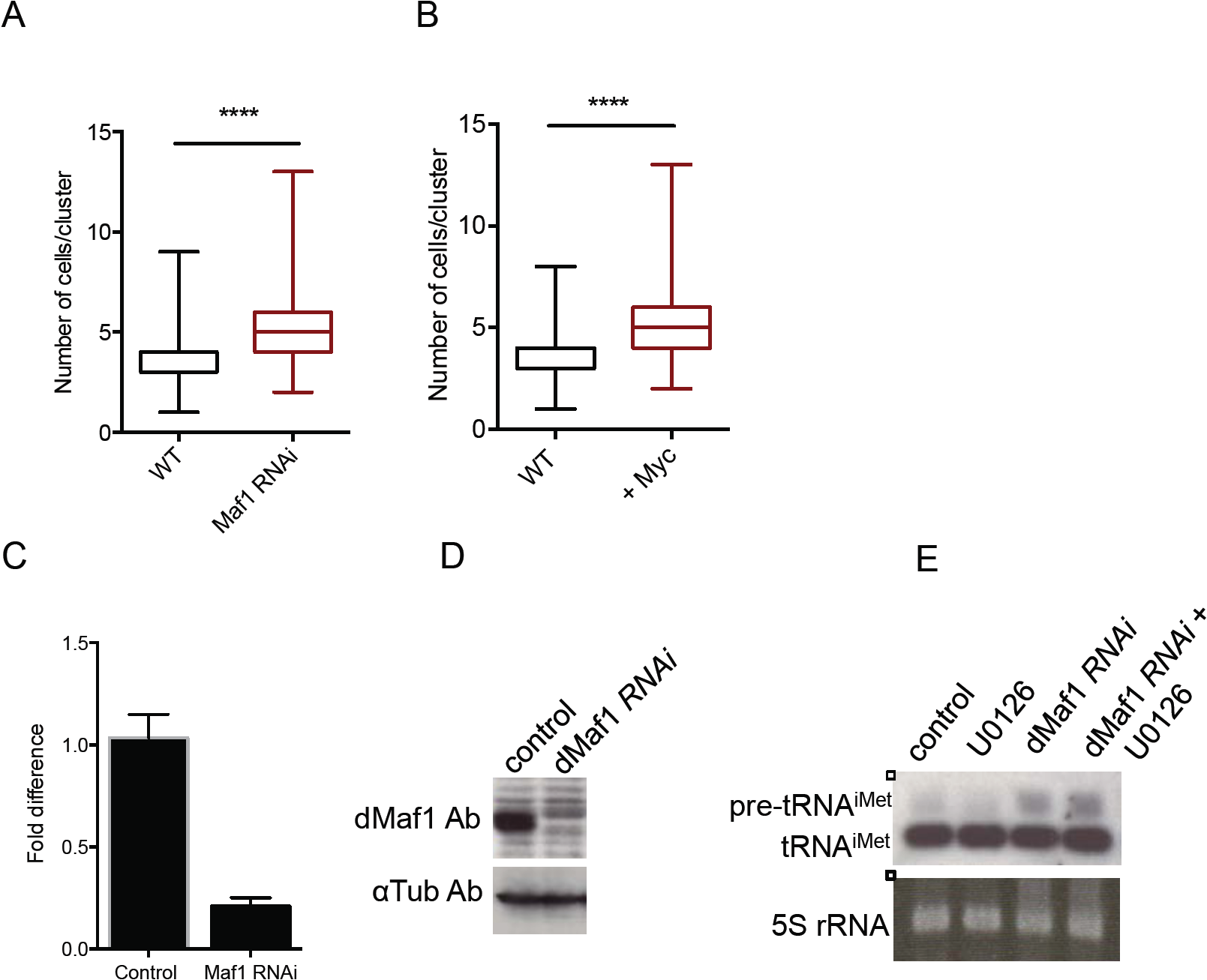
Maf1 regulates Ras-induced tRNA synthesis(related to Figure 5) (A, B) *UAS-dMaf1 RNAi* (A) or *UAS-dMyc* (B) were expressed in AMPs using the *esg-Gal4*^*ts*^ system. Larvae were shifted to 29°C at 24hrs of development and dissected at wandering stage. The numbers of cells in each AMP cluster were counted and expressed in box plots. (C) Maf1 mRNA levels were quantified by qRT-PCR data in cells treated with dsRNA to GFP (control) or dMaf1 (Maf1 RNAi) (D) Maf1 protein levels were measured by Western blot in cells treated with dsRNA to GFP (control) or dMaf1 (Maf1 RNAi). (E) dMaf1 was knocked down in *Drosophila* S2 cells by incubating cells with dsRNAs against Maf1. Control cells were treated with dsRNA to GFP. Cells were then treated with DMSO (control) or 10 μM U0126 for 2 hrs. Total RNA was isolated with Trizol and analyzed by Northern blot using DIG-labelled tRNA^iMet^ probe. Ethidium bromide stained 5S rRNA band was used as a loading control.

